# Computational Pharmacogenetics of P-Glycoprotein Mediated Antiepileptic Drug Resistance

**DOI:** 10.1101/095059

**Authors:** Ashok Palaniappan, Sindhu Varghese

## Abstract

The treatment of epilepsy using antiepileptogenic drugs is complicated by drug resistance, resulting in treatment failure in more than one-third of cases. Human P-glycoprotein (hPGP; *MDR1*) is a known epileptogenic mediator. Given that experimental investigations have suggested a role for pharmacogenetics in this treatment failure, it would be of interest to study hPGP polymorphisms that might contribute to the emergence of drug resistance. Changes in protein functional activity could result from point mutations as well as altered abundance. Bioinformatics approaches were used to assess and rank the functional impact of 20 missense MDR1 polymorphisms and the top five were selected. The structures of the wildtype and mutant hPGP were modelled based on the mouse PGP structure. Docking studies of the wildtype and mutant hPGP with four standard anti-epileptic drugs were carried out. Our results revealed that the drug binding site with respect to the wildtype protein was uniform. However the mutant hPGP proteins displayed a repertoire of binding sites with stronger binding affinities towards the drug. Our studies indicated that specific polymorphisms in MDR1 could drive conformational changes of PGP structure, facilitating altered contacts with drug-substrates and resulting in drug extrusion. This suggests that MDR1 polymorphisms could play an active role in modifying drug bioavailability, leading to pharmacoresistance in antiepileptic chemotherapy.

## 1. Introduction

P-gp (HGNC nomenclature: ABCB1) is a key transmembrane protein from bacteria to man, and it functions to protect the organism from toxic xenobiotics. P-gp has turned out to be a critical player in multiple drug resistance phenomena. Here, we are interested in its role in antiepileptic drug resistance. Epilepsy is a chronic neurological condition affecting more than 50 million people worldwide and 1-2 % of the population.1 The recurring limitation in the treatment protocol of epilepsy is the failure of drug-response in more than one-third of cases. This is the case with the *>* 30 FDA-approved drugs for epilepsy. PGP is an ATP-coupled efflux pump documented as an epileptogenic mediator.^2^ It is known to be highly expressed in the blood-brain barrier, which is pharmacologically crucial for the bioavailability of drugs acting on the central nervous system.^3^ The experimental evidence so far for the role of PGP polymorphisms in antiepileptic drug resistance has been inconclusive,^4, 5, 6, 7, 8^ but there is evidence for its cognate role in antidepressant therapy.^9^ There are at least two mechanisms by which PGP could mediate refractory epilepsy. First, elevated levels of PGP expression might be linked with the low intracellular drug concentration in cortical cells observed in epilepsy treatment. PGP is well-known for its broad substrate specificity, and would extrude drug-substrates. Alternatively, a gain-of-function mutation might enhance its functional activity, resulting in the same phenotype, i.e., hyperactive PGP leading to pharmacoresistant epilepsy.^10, 11^

PGP consists of two homologous halves, each consisting of a transmembrane (TM) domain with six alpha helices and a nucleotide-binding domain (NBD).^12^ A large, hydrophobic and polyspecific drug-binding pocket resides in the inverted V-shaped internal cavity formed by the transmembrane domains.^13^ It is clear that the key to epilepsy treatment would involve control of the epileptogenic mediator proteins, including hPGP. Structural studies of hPGP might enhance our current understanding of the role of PGP in drug resistance mechanisms, and provide any evidence of the relationship between specific *MDR1* haplotypes and altered drug pharmacokinetics. Given that little information is available on the pharmacology of missense polymorphisms of *MDR1*, analysis of the role of gain-of-function mutations in PGP would be valuable. Here, we have attempted an *in silico* study of PGP polymorphisms and their structure-activity relationships to explore drug sensitivity. PGP-mediated processes are also the major contributors to emergence of drug resistance in cancer therapy and other conditions. Our results could extend to examining the role of P-glycoprotein in generalized drug-resistance in multiple conditions.^14^

## 2. Materials and Methods

### 2.1. Polymorphism analysis

The hPGP sequence was retrieved from UniProt (acc. no. P08183). A PSI-BLAST search was performed using hPGP as query and target database as Vertebrates, with a E-value of 0.001 until convergence.15 We selected top 5000 sequences from this result, and clustered for redundancy at 40% sequence identity16. Multiple alignment of all the hits was performed using ClustalX17 and manually edited (available in Supporting Information). The dbSNP was to used to identify hPGP SNPs with the search term: “human [orgn] AND missense AND PGP”. The hits were assessed for the functional impact of polymorphisms using the curated multiple alignment obtained above. Three different tools were used: SIFT,^18^, PolyPhen2,^19^ and PhD-SNP^20^. Consensus of these predictions was used to evaluate the functionally important SNPs.

### 2.2. Homology modeling

The template structures were retrieved using a Blast search of hPGP against the PDB database.^21^ ClustalX was used to align the template and hPGP (i.e, target). Modellerv9 was used for modelling and energy-minimisation.^22^ For each target, five separate models were generated and the model with the least DOPE (discrete optimized potential energy) score was chosen as the best model. The structure of a mutant protein could be obtained by modeling in the mutation on the wild-type structure, however this would not model any global effects due to the mutation. In order to fully account for the effects of the mutation, we modelled the mutant proteins independently of the wildtype protein. Molprobity was used to validate the models obtained.^23^

**Fig. 1.**
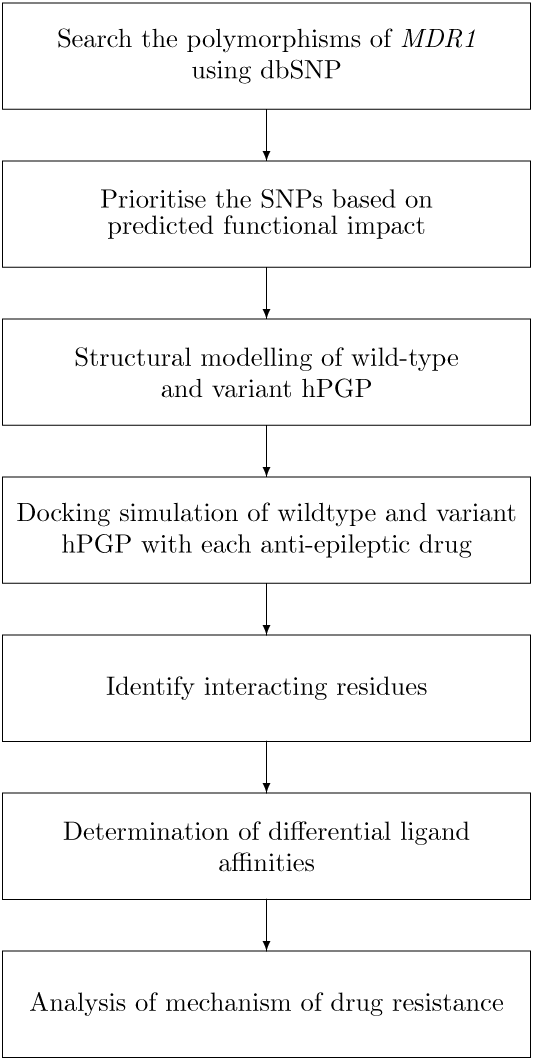
Methodology for *in silico* study of *MDR1* polymorphisms in pharmacoresistant epilepsy.

### 2.3. Protein and ligand preparation

Autodock4.2 suite of tools was used for carrying out the docking simulations of hPGP variants and anti-epileptic drugs.^24, 25^ Hydrogen atoms missing in the protein were added. This was followed by the addition of partial charges to the atoms. The protein was then converted to PDBQT format. The SMILES notation of the drugs of interest were retrieved from Pubchem.^26^ The PDB co-ordinates of the drugs of interest were generated from their SMILES representation using OpenBabel.^27^ To generate the conformers of each drug, we used MGLtools by calculating the number of bond torsions in the 3D structure. The ligand was then converted to the PDBQT format as well using AutoDock Tools. Target affinity maps for each atom type in the ligand were generated by autogrid by defining a uniform grid box centered in the hPGP internal cavity. This procedure was repeated for each target-ligand pair for a total of 6 *×* 4 = 24 times.

### 2.4. Docking

We employed the Lamarckian genetic algorithm with default parameters for docking search, with 2,500,000 cycles per run, and 10 runs per receptor-ligand pair. The binding mode with the least binding energy was defined as the best pose. The ten poses obtained for each receptor-ligand pair were clustered at 2.0 År.m.s. to validate the convergence to the best pose. The docked complex was then loaded in PDBQT format, converted to PDB coordinates using OpenBabel, and finally visualized using Rasmol2.7.28 The differential affinity of the mutant for a given ligand relative to the wildtype was estimated as the difference in the binding energies, i.e. ΔΔ*G*_*mut*_ =Δ*Gbind,mut −* Δ*Gbind,wt*.

## 3. Results and Discussion

Nearly 500 hPGP SNPs were retrieved, however most of these were unannotated, and we obtained a set of 20 hPGP SNPs for further study, none of whose functional effects were known in the literature (Table 1). The results of our assessment of functional impact by various approaches are summarised in Table 2. Most of the SNPs were determined to be neutral, not disease-causing or deleterious. Five SNPs were predicted to be functionally important by at least one of the tools, as shown in Table 2.

**Table 1.** Missense SNPs of human PGP and their location. SNPs are represented in the usual convention: wildtype aminoacid followed by position followed by replacement aminoacid.

| NO | rsID | SNP | Location |
| --- | --- | --- | --- |
| 1 | rs28381804 | F17L | N-terminal domain |
| 2 | rs9282564 | N21D | N-terminal domain |
| 3 | rs1202183 | N44S | TM1 |
| 4 | rs9282565 | A80E | Linker between TM1 and TM2 |
| 5 | *rs1128501* | G185V | TM3 |
| 6 | rs36008564 | I261V | Linker between TM4 and TM5 |
| 7 | rs2229109 | S400N | NBD1 |
| 8 | *rs28381902* | E566K | NBD1 |
| 9 | *rs28381914* | R593C | NBD1 |
| 10 | rs2235036 | A599T | NBD1 |
| 11 | rs35023033 | R669C | NBD1 |
| 12 | rs2235039 | V801M | Linker between TM8 and TM9 |
| 13 | rs2032581 | I829V | TM9 |
| 14 | rs28381967 | I836V | TM9 |
| 15 | rs2032582 | S893A | Linker between TM11 and TM12 |
| 16 | *rs72552784* | A999T | NBD2 |
| 17 | rs28401798 | P1051A | NBD2 |
| 18 | *rs55852620* | Q1107P | NBD2 |
| 19 | rs2229107 | S1141T | NBD2 |
| 20 | rs28364274 | V1251I | NBD2 |

**Table 2.** Topranked polymorphisms based on consensus prediction of functional impact

| NO | SNP | PolyPhen2<br>prediction | PolyPhen2<br>Probability | SIFT<br>prediction | SIFT<br>score | PHDSNP<br>prediction | PHDSNP<br>reliability |
| --- | --- | --- | --- | --- | --- | --- | --- |
| 1 | G185V | prob. damaging | 1 | Damaging | 1 | Disease | 8 |
| 2 | R593C | benign | 0.392 | Damaging | 1 | Disease | 6 |
| 3 | E566K | prob. damaging | 1 | Damaging | 0.88 | Disease | 6 |
| 4 | Q1107P | prob. damaging | 0.962 | tolerated | 0.19 | Disease | 6 |
| 5 | A999T | poss. damaging | 0.465 | tolerated | 1 | Neutral | 7 |

Table 3 provides the representative structures of P-glycoprotein in the PDB. Of these homologous hits, the mouse structures cover the full length of the hPGP. Some mouse structures co-crystallised with a ligand might not be representative of the native PGP conformation. When 4Q9H was superimposed with the 3G5U structure, it was observed that the register of the C-terminal half of the ‘inverted-V’ of 4Q9H was displaced relative to that of 3G5U (Fig. 3), which rendered 4Q9H unsuitable for modeling the full hPGP structure. The alignment between the hPGP and 3G5U is very good, showing *>* 87.5 % sequence identity and good sequence coverage (Fig. 2).

**Table 3.**
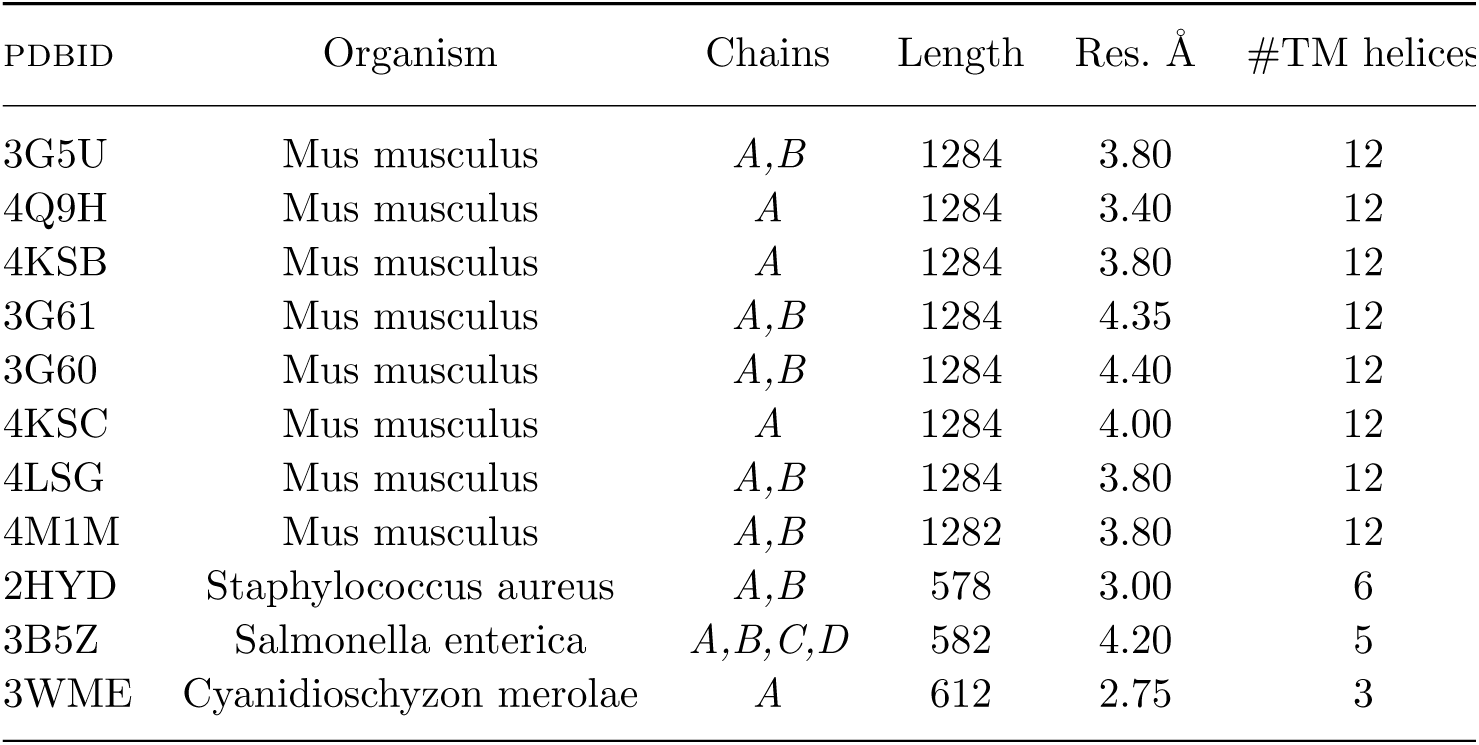
Crystal Structures of PGP homologues.

| PDBID | Organism | Chains | Length | Res. Å | #TM helices |
| --- | --- | --- | --- | --- | --- |
| 3G5U | Mus musculus | *A,B* | 1284 | 3.80 | 12 |
| 4Q9H | Mus musculus | *A* | 1284 | 3.40 | 12 |
| 4KSB | Mus musculus | *A* | 1284 | 3.80 | 12 |
| 3G61 | Mus musculus | *A,B* | 1284 | 4.35 | 12 |
| 3G60 | Mus musculus | *A,B* | 1284 | 4.40 | 12 |
| 4KSC | Mus musculus | *A* | 1284 | 4.00 | 12 |
| 4LSG | Mus musculus | *A,B* | 1284 | 3.80 | 12 |
| 4M1M | Mus musculus | *A,B* | 1282 | 3.80 | 12 |
| 2HYD | Staphylococcus aureus | *A,B* | 578 | 3.00 | 6 |
| 3B5Z | Salmonella enterica | *A,B,C,D* | 582 | 4.20 | 5 |
| 3WME | Cyanidioschyzon merolae | *A* | 612 | 2.75 | 3 |

**Fig. 2.**
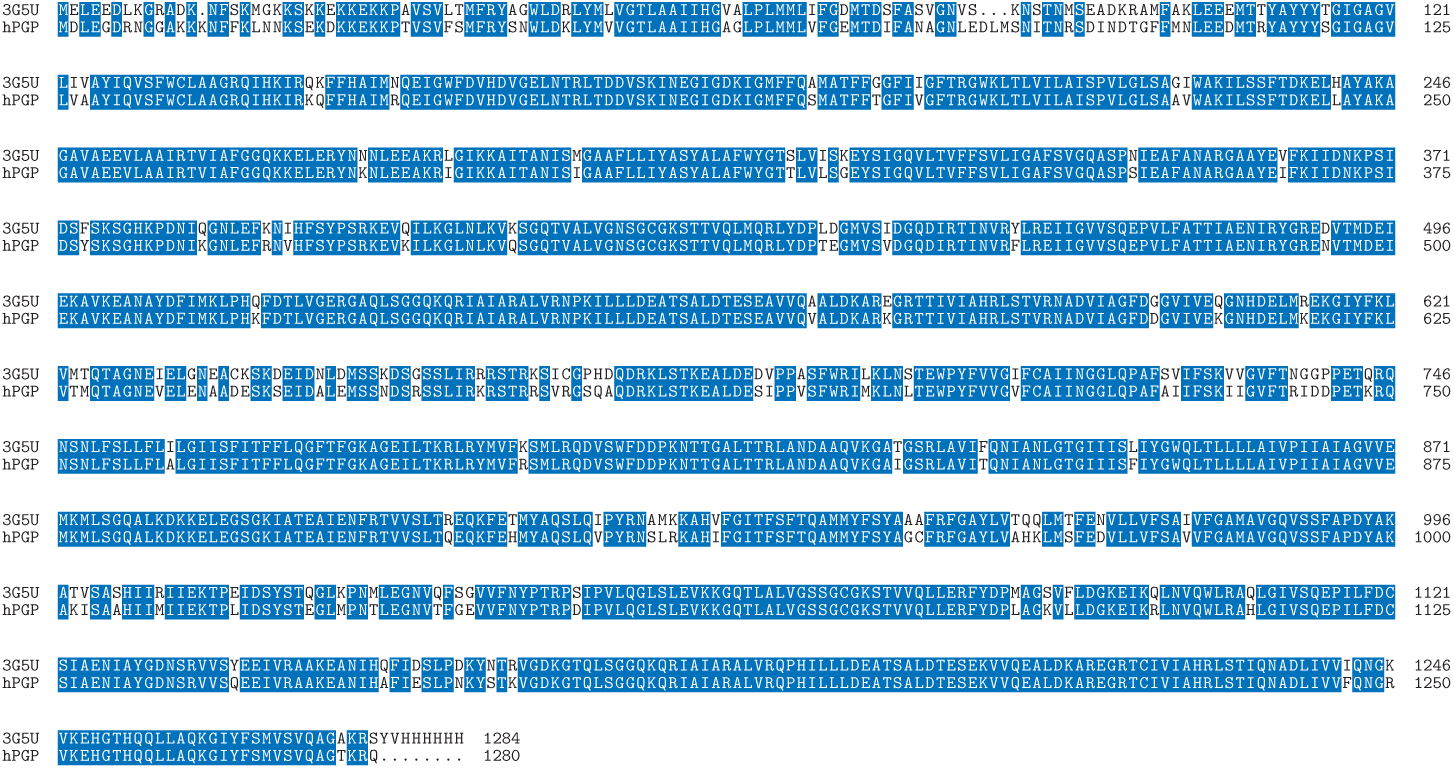
Alignment of human PGP (target) and mouse 3G5U (template). Identical residues are highlighted and gaps are indicated by..

**Fig. 3.**
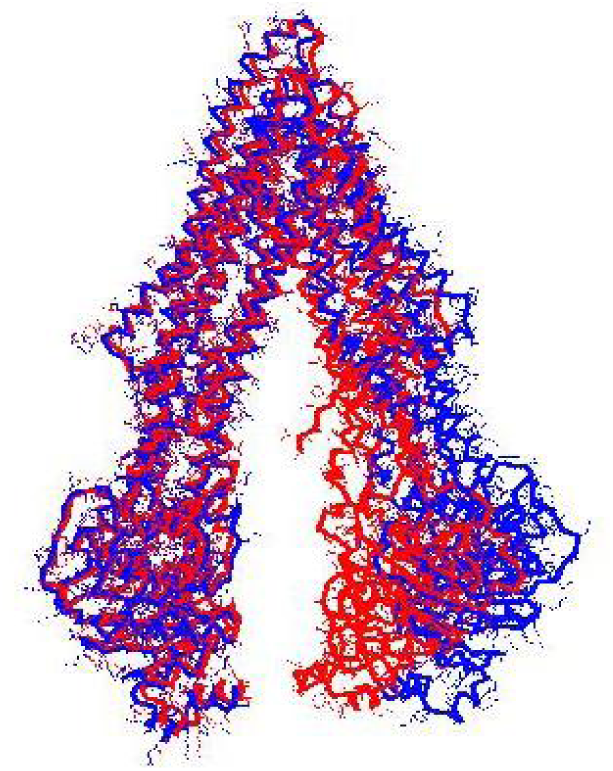
Structural superposition of 3G5U (red) and 4Q9H (blue). Note the displacement of the C-terminal region of 4Q9H.

3G5U was used as the template for homology modeling. The target structures of the hPGP wildtype and the five variants were independently modeled and energy minimised five times each, and the best model was used for further studies. All the modelled structures are available in the Supporting Information and their mutual rms deviations are shown in Table 4.

**Table 4.** Mutual rms deviation of the template and modelled structures (in Å). Mutant numbering corresponds to the order in Table 2.

|  | 3g5u | wt | Mut1 | Mut2 | Mut3 | Mut4 |
| --- | --- | --- | --- | --- | --- | --- |
| wt | 4.51 |  |  |  |  |  |
| Mut1 | 4.24 | 0.53 |  |  |  |  |
| Mut2 | 4.43 | 0.62 | 1.31 |  |  |  |
| Mut3 | 4.32 | 0.71 | 1.19 | 1.33 |  |  |
| Mut4 | 4.33 | 0.43 | 0.97 | 0.81 | 0.59 |  |
| Mut5 | 4.63 | 0.49 | 0.71 | 1.04 | 0.48 | 0.44 |

Phenobarbital was first used as an antiseizure drug in 1912, followed by phenytoin. Today more than 30 drugs are FDA-approved in the treatment of epilepsy, yet all of them face pharmacoresistance and more than one-third of epilepsy cases remain untreatable. In addition to phenobarbital and phenytoin, two other common antiepileptic medications, namely valproate and carbamazepine, were included in the set of ligands studied (Table 5).

**Table 5.** Anti-epileptic drugs.

| NO | Drug | PUBCHEMID | SMILES notation |
| --- | --- | --- | --- |
| 1 | Valproate | 3121 | CCCC(CCC)C(=O)O |
| 2 | Phenytoin | 1775 | C1=CC=C(C(=C1)C2(C(=O)NC(=O)N2)C3=CC=CC=C3 |
| 3 | Carbamazepine | 2554 | C1=CC=C2C(=C1)C=CC3=CC=CC=C3N2C(=O)N |
| 4 | Phenobarbital | 4763 | CCC1(C(=O)NC(=O)NC1=O)C2=CC=CC=C2 |

Docking between each hPGP protein (wildtype + 5 mutants) and ligand was carried out. Ten docking runs were performed per receptor-ligand pair. Each run provides one low-energy docked conformation of the respective receptor-ligand pair.

The site corresponding to the lesat-energy binding mode was taken as the binding site of the ligand with the receptor. To ascertain convergence to the lowest-energy binding mode, the ten runs of each receptor-ligand binding conformations were clustered at 2.0Å r.m.s. The lowest-energy binding modes showed good convergence. Of the 24 receptor-ligand pairs, 21 had energy-histograms showing the least binding energy (*±*0.2 *kcal/mol*) as the most probable conformation and the least binding energies of the rest were within 1 *kcal/mol* of the binding energies of the most probable conformation. This provided confidence that the docking procedure resulted in convergence to the optimum receptor-ligand conformation. The structures of the receptor-drug complexes as well as the best poses (defined as within 4.5Å of the ligand) are available in the Supporting Information. A comparison of the best poses between the wildtype and one of the mutants is illustrated in Fig. 4.

**Fig. 4.**
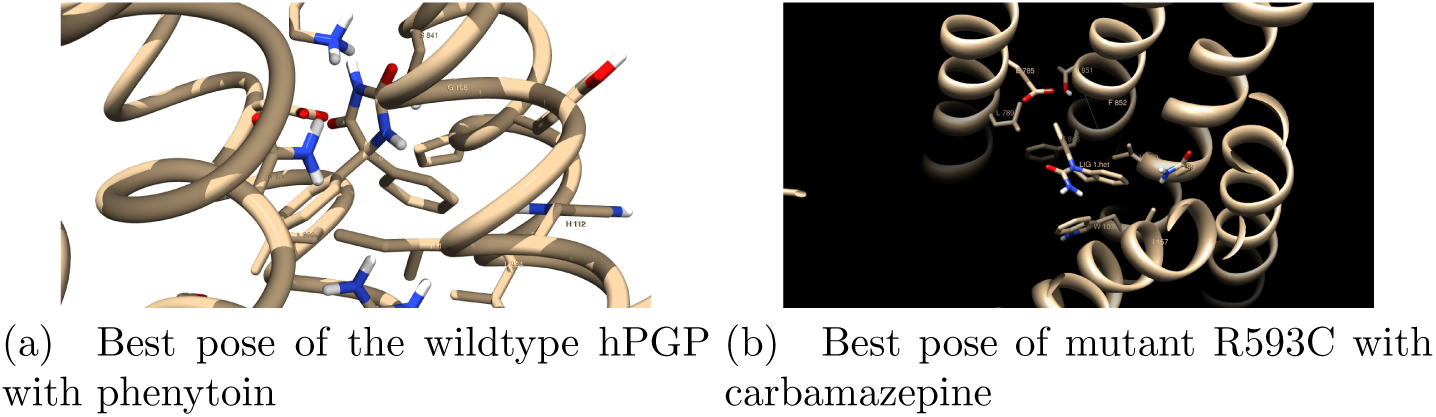
Illustration of best pose comparisons.

The hPGP residues binding the ligand in each hPGP-drug pair represent the drug-specific binding pocket. These residues were defined at a contact distance of *<* 4.5Å from the drug in the bound conformation. These residues contributed to stabilizing the docked complex by forming hydrogen bonds and Van der Waals interactions with the substrates. The groups of contacting residues specific to each docked complex are shown in Table 6. The conservation of these residues calculated using the constructed multiple alignment is given in the Supporting information. It is observed that the binding site of wildtype hPGP is identical for phenytoin and carbamazepine. This is a 14-residue binding pocket in the internal cavity lined by four hydrophobic residues (Ile144, Val179, Leu890, Leu924), three charged residues (Arg148, Asp886, Lys934) and four polar residues (Ser180, Asn183, Tyr928, Ser931). In contrast, the mutant proteins bound the drugs in alernative variable regions, notably a binding pocket involving Gln99, Val100, Trp103, Ile157, Glu785, Leu789, Phe848, Thr851, and Phe852 that bound all drugs except valproate. It was remarkable that for a given mutant hPGP, the binding pocket interactions differed for each drug. The R593C hPGP mutant bound phenytoin very close to the mutation site, suggesting evidence for local conformational change in binding the drug.

**Table 6.**
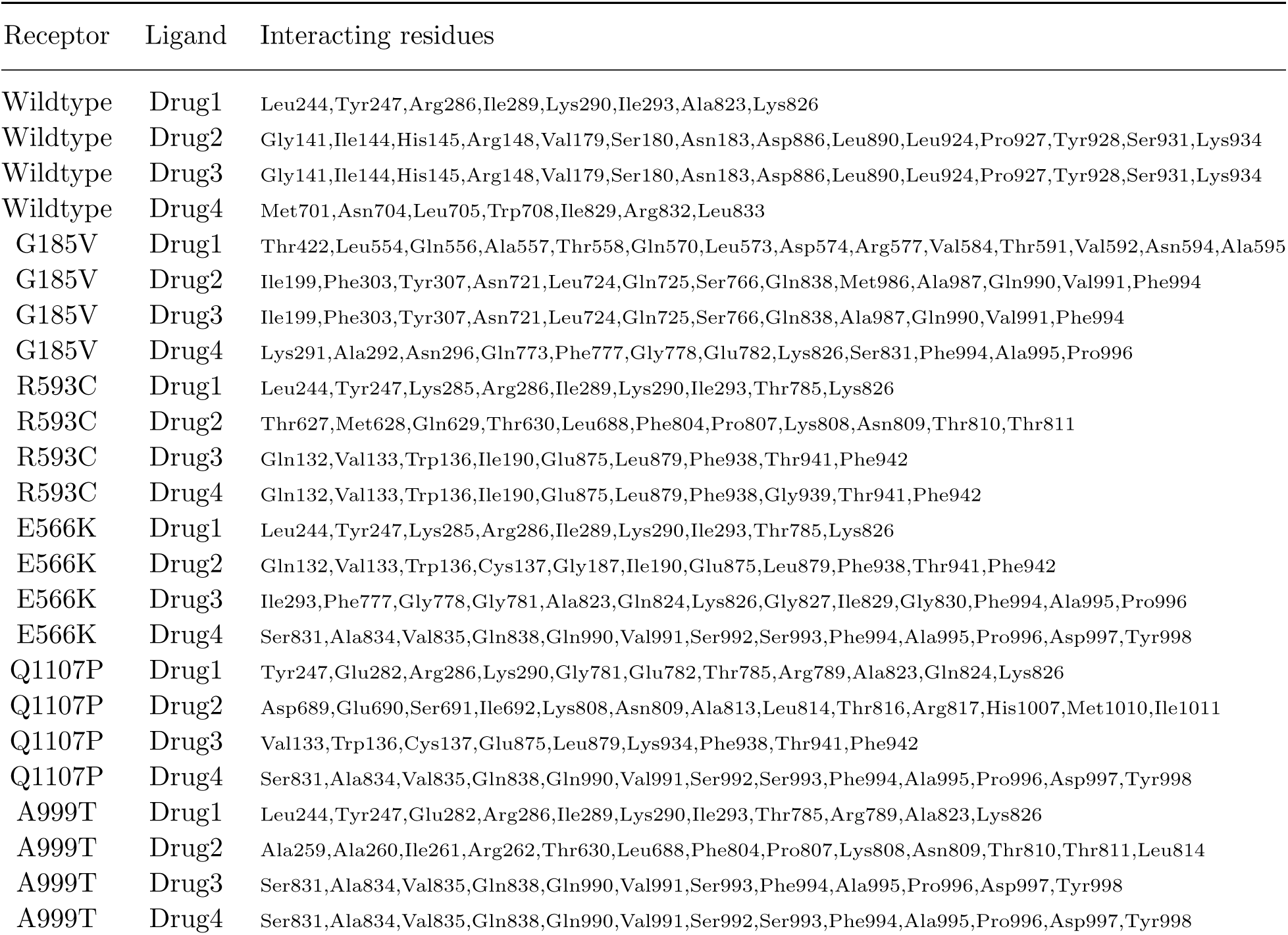
Contacting residues of the receptor within 4.5° Åof the ligand in the best pose of each docked ligand-receptor pair. Residue numbering follows the UniProt human P-gp entry P08183.

| Receptor | Ligand | Interacting residues |
| --- | --- | --- |
| Wildtype | Drug1 | Leu244,Tyr247,Arg286,Ile289,Lys290,Ile293,Ala823,Lys826 |
| Wildtype | Drug2 | Gly141,Ile144,His145,Arg148,Val179,Ser180,Asn183,Asp886,Leu890,Leu924,Pro927,Tyr928,Ser931,Lys934 |
| Wildtype | Drug3 | Gly141,Ile144,His145,Arg148,Val179,Ser180,Asn183,Asp886,Leu890,Leu924,Pro927,Tyr928,Ser931,Lys934 |
| Wildtype | Drug4 | Met701,Asn704,Leu705,Trp708,Ile829,Arg832,Leu833 |
| G185V | Drug1 | Thr422,Leu554,Gln556,Ala557,Thr558,Gln570,Leu573,Asp574,Arg577,Val584,Thr591,Val592,Asn594,Ala595 |
| G185V | Drug2 | Ile199,Phe303,Tyr307,Asn721,Leu724,Gln725,Ser766,Gln838,Met986,Ala987,Gln990,Val991,Phe994 |
| G185V | Drug3 | Ile199,Phe303,Tyr307,Asn721,Leu724,Gln725,Ser766,Gln838,Ala987,Gln990,Val991,Phe994 |
| G185V | Drug4 | Lys291,Ala292,Asn296,Gln773,Phe777,Gly778,Glu782,Lys826,Ser831,Phe994,Ala995,Pro996 |
| R593C | Drug1 | Leu244,Tyr247,Lys285,Arg286,Ile289,Lys290,Ile293,Thr785,Lys826 |
| R593C | Drug2 | Thr627,Met628,Gln629,Thr630,Leu688,Phe804,Pro807,Lys808,Asn809,Thr810,Thr811 |
| R593C | Drug3 | Gln132,Val133,Trp136,Ile190,Glu875,Leu879,Phe938,Thr941,Phe942 |
| R593C | Drug4 | Gln132,Val133,Trp136,Ile190,Glu875,Leu879,Phe938,Gly939,Thr941,Phe942 |
| E566K | Drug1 | Leu244,Tyr247,Lys285,Arg286,Ile289,Lys290,Ile293,Thr785,Lys826 |
| E566K | Drug2 | Gln132,Val133,Trp136,Cys137,Gly187,Ile190,Glu875,Leu879,Phe938,Thr941,Phe942 |
| E566K | Drug3 | Ile293,Phe777,Gly778,Gly781,Ala823,Gln824,Lys826,Gly827,Ile829,Gly830,Phe994,Ala995,Pro996 |
| E566K | Drug4 | Ser831,Ala834,Val835,Gln838,Gln990,Val991,Ser992,Ser993,Phe994,Ala995,Pro996,Asp997,Tyr998 |
| Q1107P | Drug1 | Tyr247,Glu282,Arg286,Lys290,Gly781,Glu782,Thr785,Arg789,Ala823,Gln824,Lys826 |
| Q1107P | Drug2 | Asp689,Glu690,Ser691,Ile692,Lys808,Asn809,Ala813,Leu814,Thr816,Arg817,His1007,Met1010,Ile1011 |
| Q1107P | Drug3 | Val133,Trp136,Cys137,Glu875,Leu879,Lys934,Phe938,Thr941,Phe942 |
| Q1107P | Drug4 | Ser831,Ala834,Val835,Gln838,Gln990,Val991,Ser992,Ser993,Phe994,Ala995,Pro996,Asp997,Tyr998 |
| A999T | Drug1 | Leu244,Tyr247,Glu282,Arg286,Ile289,Lys290,Ile293,Thr785,Arg789,Ala823,Lys826 |
| A999T | Drug2 | Ala259,Ala260,Ile261,Arg262,Thr630,Leu688,Phe804,Pro807,Lys808,Asn809,Thr810,Thr811,Leu814 |
| A999T | Drug3 | Ser831,Ala834,Val835,Gln838,Gln990,Val991,Ser993,Phe994,Ala995,Pro996,Asp997,Tyr998 |
| A999T | Drug4 | Ser831,Ala834,Val835,Gln838,Gln990,Val991,Ser992,Ser993,Phe994,Ala995,Pro996,Asp997,Tyr998 |

Further clarity on these observations could be obtained on an examination of the estimated binding free energies of the wildtype and variant hPGPs with the different drugs of interest. Table 7 shows these binding energies along with the predicted differential ligand affinity which is estimated by ΔΔ*G*_*mut*_ = Δ*G*_*bind,mut*_*−*Δ*G*_*bind,wt*_. It was observed that all but three of the differential ligand affinities were negative. This implied that the variant hPGP bound each drug with a stronger affinity than the wildtype hPGP. The maximum range of differential response was observed with mutant E566K (*−*1.25 *kcal/mol <* ΔΔ*G*_*mut*_ < 0.80 *kcal/mol*).

**Table 7.** Free energies of binding (Δ*G*_*bind*_) of each docked receptor-drug pair. The predicted differential ligand affinity is given by ΔΔ*G*_*mut*_ = Δ*G*_*bind,mut*_ *−*Δ*G*_*bind,wt*_. All values in *kcal/mol*.

| Receptor | valproate | phenytoin | carbamazepine | phenobarbital |
| --- | --- | --- | --- | --- |
| wildtype | -4.28 | -5.48 | -6.35 | -5.24 |
| G185V | -3.90 | -6.38 | -6.27 | -5.62 |
| R593C | -5.21 | -5.52 | -6.37 | -5.74 |
| E566K | -5.53 | -6.17 | -5.55 | -6.15 |
| Q1107P | -5.50 | -5.47 | -6.66 | -5.52 |
| A999T | -5.15 | -5.65 | -6.67 | -6.48 |
| $\Delta\Delta G_{G185V}$ | 0.38 | -0.90 | 0.08 | -0.38 |
| $\Delta\Delta G_{R593C}$ | -0.93 | -0.04 | -0.02 | -0.5 |
| $\Delta\Delta G_{E566K}$ | -1.25 | -0.69 | 0.80 | -0.91 |
| $\Delta\Delta G_{Q1107P}$ | -1.22 | 0.01 | -0.31 | -0.28 |
| $\Delta\Delta G_{A999T}$ | -0.87 | -0.17 | -0.32 | -1.24 |

Two features indicated the neutrality of wildtype hPGP with respect to binding anti-epileptic drugs. First, the binding pocket appeared constant for both phenytoin and carbamazepine. Second, the binding energy with the drug was higher relative to the variants and hence less tight. On the other hand, there were two features that indicated that mutant hPGPs would assist in the development of drug resistance. First, mutant hPGPs bound each drug in a different location in the internal cavity. Variability in location affords a better search of the optimal binding modes of the drug. Second, consistently lower binding energies were observed, implying stable drug-PGP complexes for possible energetic extrusion of the drug. The *in silico* analysis showed that polymorphisms could have played a role in relocating the optimal drug-binding cavity for a higher affinity, relative to the wild-type hPGP.

An elevated affinity between a mutant hPGP and the drug could suggest a potential differential adverse response to therapy. From Table 7, it is seen that this is the case for 17 out of the 20 drug-protein combinations studied. Experimental studies are necessary to validate these results and determine whether the magnitude of any differential adverse responses could translate to the threshold for the development of pharmacoresistance.

## 4. Conclusions

Though hPGP is well-documented as a modifier of drug bioavailability in many conditions, its role in antiepiletic drug resistance has been controversial. At least two alternative mechanisms could explain the hPGP-mediated epileptogenic phenotype.

Our work suggests that polymorphisms are a viable mechanism of PGP action that could lead to drug resistance acquisition independent of other mechanisms. It is interesting that all the polymorphisms appeared to result in gain-of-function. Coupled with the observation that somatic mutations could have a similar effect to identical inherited polymorphisms, this would suggest that PGP is a potential oncogene in the context of cancer drug resistance.

Developing a drug resistance strategy to combat drug resistance is a top priority. Our work has highlighted that *MDR1* polymorphisms could potentially lower the threshold for development of pharmacoresistance. This gain-of-function process in hPGP offers a novel candidate target in the fight against antiepileptic drug resistance. Experimental validation of our work is necessary to apply our findings towards achieving pharmacosensitive response in epilepsy treatment. Our methodology is extendable to studies investigating the effect of genetic polymorphisms on phenotypes in other diseases and conditions.

## 5. Supporting Information

Supporting information is available at 10.6084/m9.figshare.5937388.

